# PplT domain proteins – ubiquitous potential prokaryotic phospholipid translocases

**DOI:** 10.1101/2022.03.11.483950

**Authors:** Janna N. Hauser, Arnaud Kengmo Tchoupa, Susanne Zabel, Kay Nieselt, Christoph M. Ernst, Christoph J. Slavetinsky, Andreas Peschel

## Abstract

Flippase proteins exchanging phospholipids between the cytoplasmic membrane leaflets have been identified in Eukaryotes but remained largely unknown in Prokaryotes. Only MprF proteins that synthesize aminoacyl phospholipids in some bacteria have been found to contain a domain that translocates the produced lipids. We show here that this domain, which we named ‘prokaryotic phospholipid translocator’ (PplT), is widespread in Bacteria and Archaea, encoded as a separate protein or fused to different types of enzymes. We also demonstrate that the *Escherichia coli* PplT protein interacts with many phospholipid-synthetic enzymes and deletion of *pplT* impaired bacterial growth, which supports its potential role in membrane lipid metabolism. PplT domain proteins may be general prokaryotic lipid flippases with critical roles in cellular homeostasis.

## Introduction

Most of the basic processes and enzyme systems involved in growth and maintenance of prokaryotic cells have been explored in the past. Major mechanisms of membrane lipid synthesis and turnover have also been described thoroughly [1, 2]. However, ‘flippase’ proteins that facilitate the translocation of phospholipids from the inner leaflet, the site of synthesis, to the outer leaflet of the cytoplasmic membrane, have remained largely unknown. We speculated that such flippases might be among the conserved membrane proteins of unknown function found in many prokaryotic taxa. Phospholipid flippases may not be essential for cell viability, which might explain why they have not been identified in previous screening approaches for prokaryotic proteins with crucial function for cellular integrity. Flippases have been characterized in eukaryotic cells [3] but the responsible proteins do not seem to have obvious homologs in prokaryotes.

Flippase reactions are particularly difficult to study because the flipping process is fast and hard to monitor as the lipids remain in place and adapt only their orientation. Nevertheless, a prototype flippase has been identified in prokaryotic MprF proteins that synthesize and then translocate aminoacyl phosphatidylglycerol (Aa-PG) lipids such as lysyl-phosphatidylglycerol (Lys-PG) or alanyl-phosphatidylglycerol (Ala-PG) [4, 5]. Those lipids reduce the negative net charge at the outer surface of cytoplasmic membranes, thereby diminishing the binding of cationic antimicrobial molecules such as bacterial bacteriocin and host defensin peptides [6]. To achieve protection against such molecules, it is critical for microorganisms to efficiently translocate Lys-PG or Ala-PG, once these lipids have been formed by the MprF synthase domain at the inner surface of the cytoplasmic membrane. The synthase domain cooperates with the flippase domain in a dynamic fashion to deliver the lipid products to the translocation channel [7]. The two domains function together even if expressed as separate proteins [8]. The flippase domain of MprF consists of eight transmembrane sections [8] and contains a motif that is accessible from both sides of the cytoplasmic membrane [9]. It has been proposed that this part moves during the lipid translocation process or that it is in the center between two larger cavities that form the lipid translocation channel [9]. The structure of the *Rhizobium tropici* MprF protein with bound Lys-PG molecules captured in the process of translocation has recently been elucidated by cryoelectron microscopy [7]. It gives rise to a mechanistic model that proposes a process with Lys-PG entering an inner protein cavity, which transiently opens a gate to an outer cavity, from which Lys-PG is subsequent released to the outer membrane leaflet [7]. The extent to which the flippase domain found in MprF proteins is present in Bacteria and Archaea, and if it may play a more general role in phospholipid translocation, has remained unknown.

The flippase domain of MprF proteins exhibits a conserved architecture with low sequence identity. It is referred to as cl04219, pfam03706, or UPF0104 in the NCBI Conserved Domain Database (CDD) [10] and as IPR022791 in the InterPro protein signatures database [11]. We show here that proteins consisting of this domain alone or in combination with various enzymatic domains are found in almost all of the major groups of Bacteria and Archaea, probably to accomplish the translocation of membrane lipids. The corresponding *Escherichia coli* protein interacts with most of the phospholipid-biosynthetic enzymes and it is encoded in the vicinity of a cardiolipin (CL) synthase. Its inactivation is not lethal but affects the fitness of *E. coli*. We suggest naming the corresponding flippase domain ‘prokaryotic phospholipid translocator’ (PplT), which is likely to fulfill a general membrane homeostatic function in prokaryotes.

## Results

### 1. PplT domain proteins are widespread in most groups of prokaryotes

MprF-related proteins are found in many different bacterial taxa [12]. Notably, we found that the two functional domains of MprF occur in different combinations. The Lys-PG synthase domain, for instance, occurs sometimes without the Lys-PG flippase domain [12]. On the other hand, the flippase domain is often encoded without a Lys-PG synthase or in combination with other enzymatic domains. Systematic evaluation of microbial genomes for the presence of this domain, referred to as PplT (cl04219), revealed an almost universal occurrence in most phyla of Bacteria (Fig. 1A) and in several Classes of Archaea (Fig. 1B). Indeed, blastp searches uncovered that the PplT domain is even found in bacterial species, which do not produce Aa-PGs, such as *E. coli* and many other Enterobacteriaceae [13, 14]. Thus, PplT may have a broader role in prokaryotes than previously thought. It should be noted that most prokaryotes with a *pplT* homolog encode only one protein with a PplT domain in their genome (Fig. S1).

**Figure 1:**
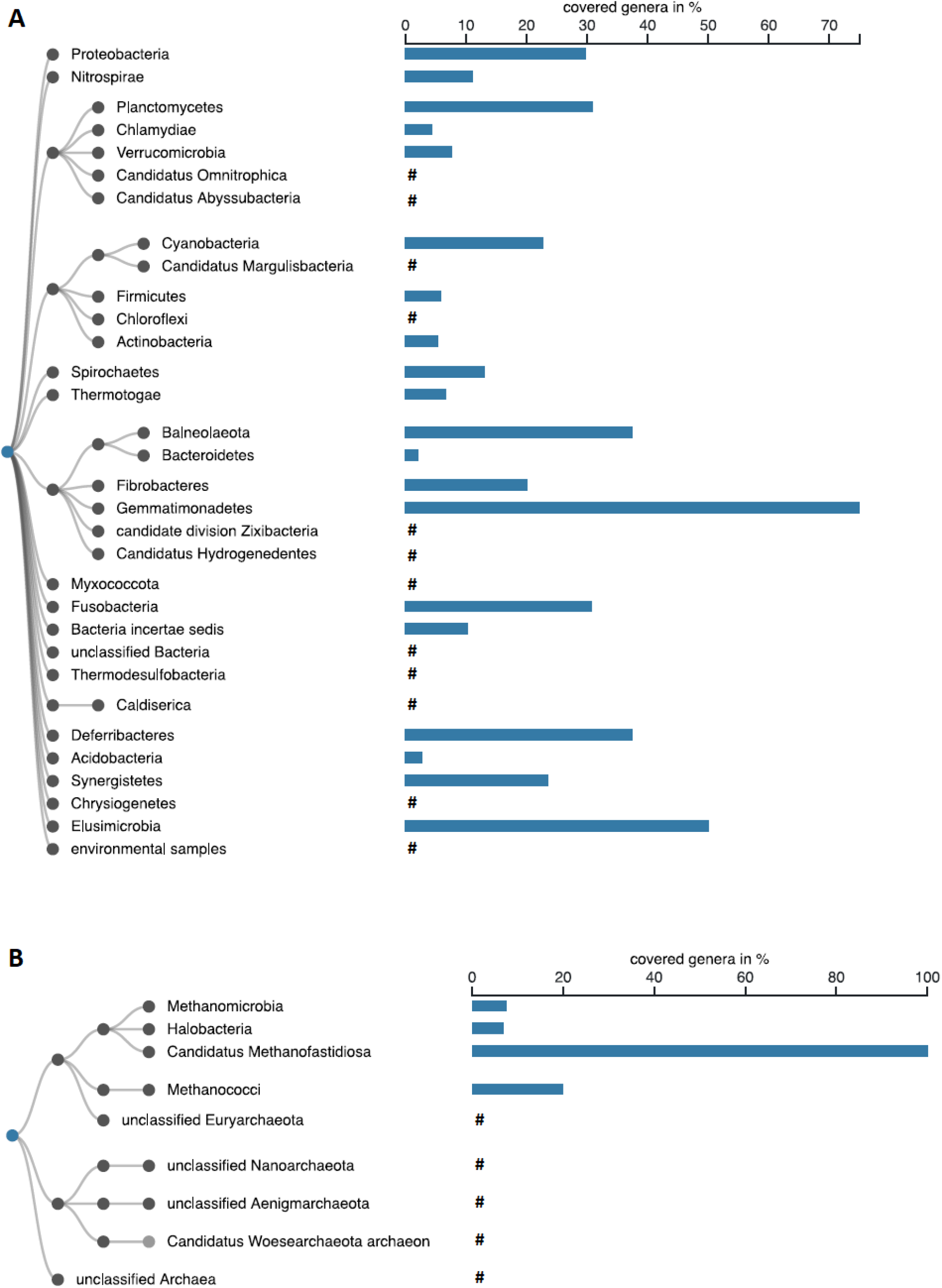
Distribution of the PpIT domain amongst taxa of the Bacteria (A) and Archaea (B) superkingdoms. The taxonomy was collapsed to the phylum rank. Dark grey leaf nodes indicate collapsed nodes. The bar chart indicates the fraction of genera of a particular Phylum (Bacteria) or Class (Archaea) carrying a PpIT homolog within the subtree rooted at the respective taxon. #, taxa include *pplT* genes but percentages of covered genera could not be calculated due to incomplete assignment of the genome on genus level.

PplT occurs as a separate protein in many different bacterial phyla including Proteobacteria, Actinobacteria, Firmicutes, Bacteroidetes, Planctomycetes, and Thermotogae as well as archaeal phyla including Euryarchaeota and Crenarchaeota. Moreover, PplT is found connected to other types of proteins in addition to Lys-PG or Ala-PG synthases (Fig. 2). The Conserved Domain Architecture Retrieval Tool [15] lists 148 different architectures of PplT domain-containing proteins. The most prevalent architectures include combinations with Aa-PG synthase domain (cl41273) of MprF proteins, glycosyl transferase domains (family A, cl11394; or family B, cl10013), or protein kinase domains (cl21453) (Fig. 2). MprF proteins were most often found in genomes of Cyanobacteria, Actinobacteria, Firmicutes, and Proteobacteria. A C-terminal or N-terminal family-B glycosyltransferase domain was found attached to PplT proteins from Actinobacteria, Nitrospirae, and Chloroflexi or Acidobacteria, respectively, while a family-A glycosyltransferase domain was found at the N-terminus of PplT in Archaea of the Euryarchaeota phylum. Several Actinobacteria encode a protein kinase domain, N-terminally fused to PplT. Further, less frequently occurring combinations were found with enzymatic domains such as dehalogenase-like hydrolases (cl21460), O-antigen ligases (cl04850), S-adenosylmethionine-dependent methyltransferase (cl17173), or phosphotransferases (cl37506), to name a few (Fig. 2). These findings suggest that PplT-containing proteins may have multiple roles, presumably in the synthesis, modification, and translocation of different types of membrane lipids or in membrane-linked signal transduction processes. In several bacterial and archaeal taxa, only a minority of the genera appear to encode PplT proteins in their genomes (Fig. 1) suggesting that PplT proteins may be important only in certain habitats or that some bacteria may use other transport systems, that could translocate phospholipids along with other types of cargo molecules.

**Figure 2:**
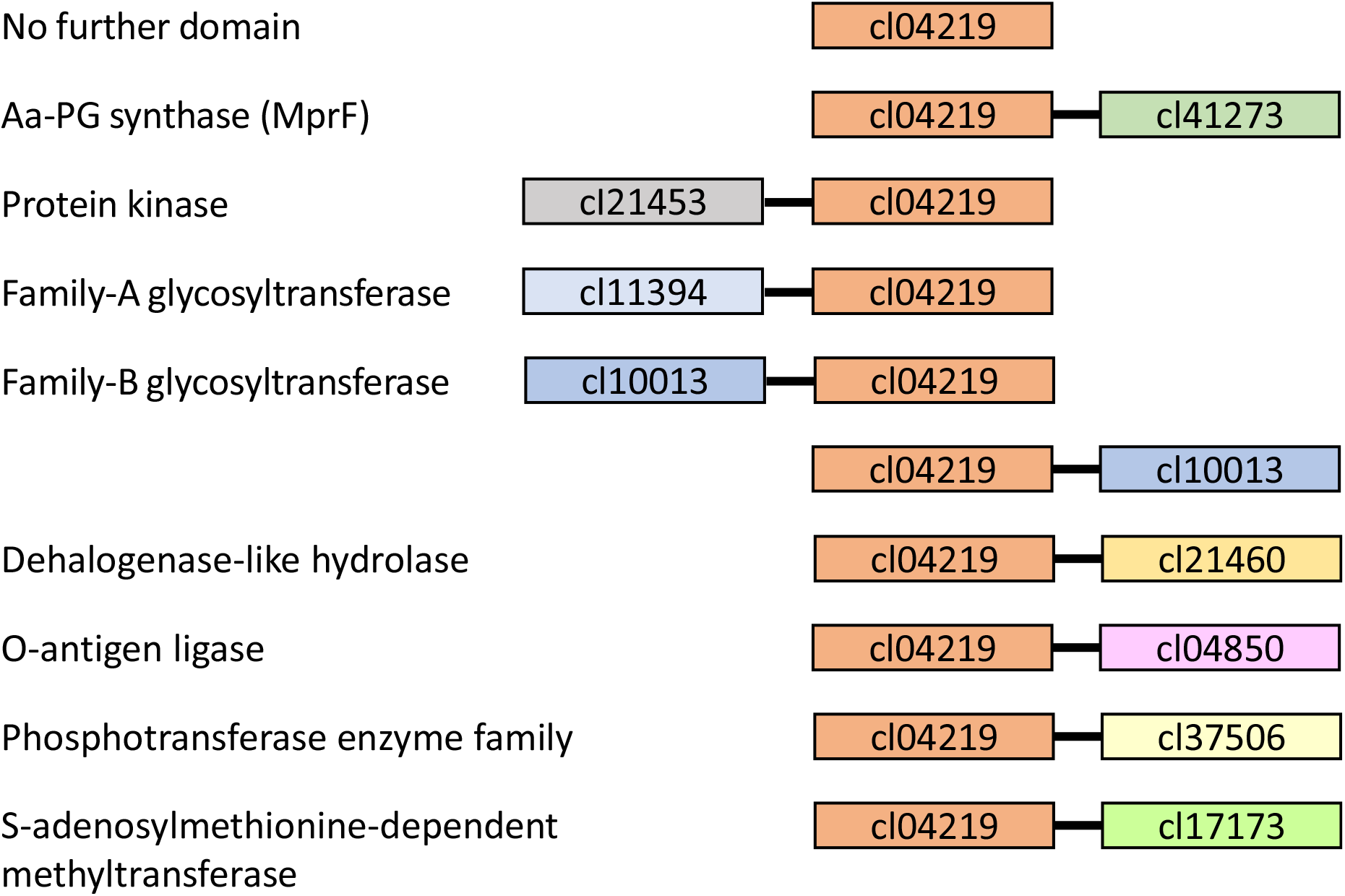
PplT domain protein architectures. Combinations of the PplT domain (cI04219) with other protein domains found in Bacteria and Archaea. The 10 most commonly occurring domain combinations are shown, as revealed by the Conserved Domain Architecture Retrieval Tool [15]. The domains are not drawn to scale.

### 2. The *E. coli* PplT is encoded together and interacts with lipid-synthetic enzymes

In order to study potential functions of PplT proteins not linked to other protein domains, we focused on the *E. coli* homolog, a protein of unknown function with the generic name YbhN, we suggest to rename PplT. *pplT* is not essential in *E. coli,* mutations in this gene are included in the Keio collection, a transposon mutant library [16], albeit without a reported phenotype. *pplT* is encoded in an operon together with the CL synthase gene *clsB* in *E. coli* and other Enterobacteriaceae including the genera *Shigella, Klebsiella* and *Pantoea* (Fig. 3A), which is in agreement with the role of PplT in lipid synthesis and translocation.

**Figure 3:**
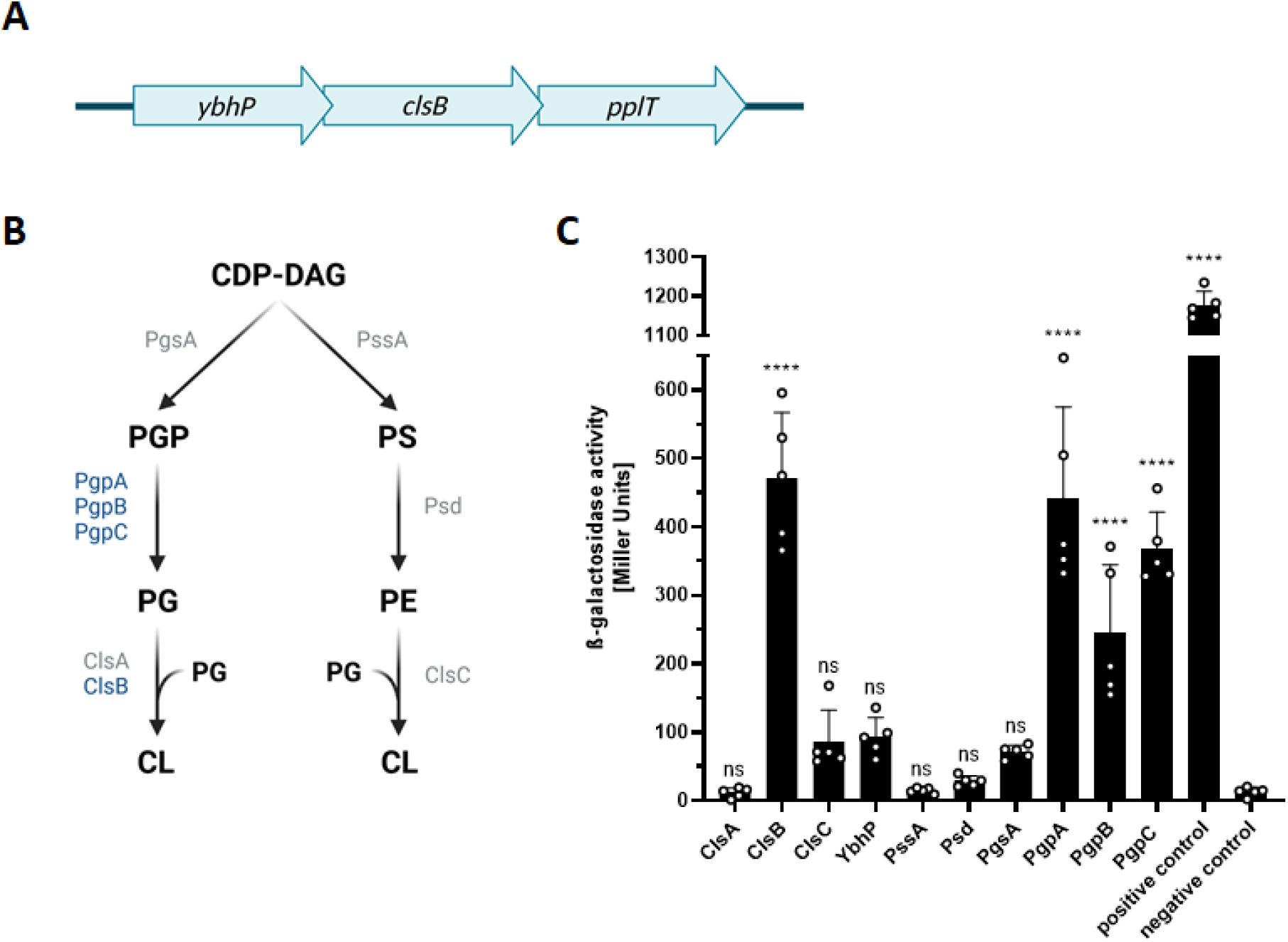
Genetic context and interaction partners of PplT in *E. coli*. (A) PplT is encoded in an operon together with YbhP, a protein of unknown function, and the cardiolipin synthase ClsB in Enterobacteriaceae, including the genera *Escherichia, Shigella*, *Klebsiella,* and *Pantoea*. The genes of the operon are not drawn to scale. (B) Schematic representation of the phospholipid synthesis pathways in *E. coli*. Cytidine diphosphate diacylglycerol (CDP-DAG), phosphatidylglycerol-3-phosphate (PGP), phosphatidylglycerol (PG), cardiolipin (CL), phosphatidylserine (PS), phosphatidylethanolamine (PE). Enzymes are marked in blue or grey, if they interacted with PplT or not, respectively. (C) Interaction studies with PplT and lipid-biosynthetic enzymes using the bacterial two-hybrid system and β-galactosidase assay for quantification. Data represent means plus SD of five biological replicates. Statistical significances were determined by two-way ANOVA followed by Dunnett’s test in comparison to the negative control (ns, not significant; ****, P <0.0001). Figures A and B were created with BioRender.com.

Phospholipid-biosynthetic proteins have been found to form large multicomponent complexes in bacteria such as *S. aureus* [17, 18]. To analyze a role of PplT in such complexes, its potential interaction with several phospholipid-biosynthetic proteins was studied in the bacterial two-hybrid system [8, 19, 20]. The genes to be analyzed were cloned in *E. coli* expression plasmids, resulting in C- and N-terminal fusions of potential interaction partners or PplT with the complementary adenylate cyclase fragment T18 or T25, respectively. Interaction of T18 with T25 results in the production of cAMP, leading to expression of β-galactosidase, which was quantified. *E. coli* does not produce aminoacyl phospholipids but, in addition to phosphatidylglycerol (PG) and CL, phosphatidylserine (PS) and phosphatidylethanolamine (PE) [2]. PplT was found to interact with the proteins PgpA, PgpB, and PgpC, involved in PG synthesis and with the CL synthase ClsB, encoded in the same operon as PplT (Fig. 3A, B, C). Hence, PplT interacts with the majority of PG and CL biosynthetic enzymes in *E. coli* (Fig. 3C), which supports a role of PplT in phospholipid metabolism.

### 3. PplT affects *E. coli* fitness

Although *pplT* is not essential in *E. coli,* its ubiquitous presence in most of the prokaryotic taxa suggests an important role in basic cellular functions. *pplT* was deleted in the uropathogenic *E. coli* (UPEC) strain CFT073. The mutant had the same phospholipid pattern as the wild type (WT) indicating that PplT has no obvious impact on the overall lipid composition (Fig. 4A, B). The *pplT* mutant exhibited a growth defect compared with the WT in the late logarithmic phase and it did not reach the cell density of WT cultures (Fig. 5). This phenotype was pronounced at 37°C (Fig. 5A) but not observed at 42°C (Fig. 5B), which supports a role of PplT in optimal membrane function, a process that depends on temperature. When co-cultivated over several days, the WT grew better than the *pplT* mutant (Fig. 5C, D), demonstrating a competitive fitness benefit conferred by PplT.

**Figure 4:**
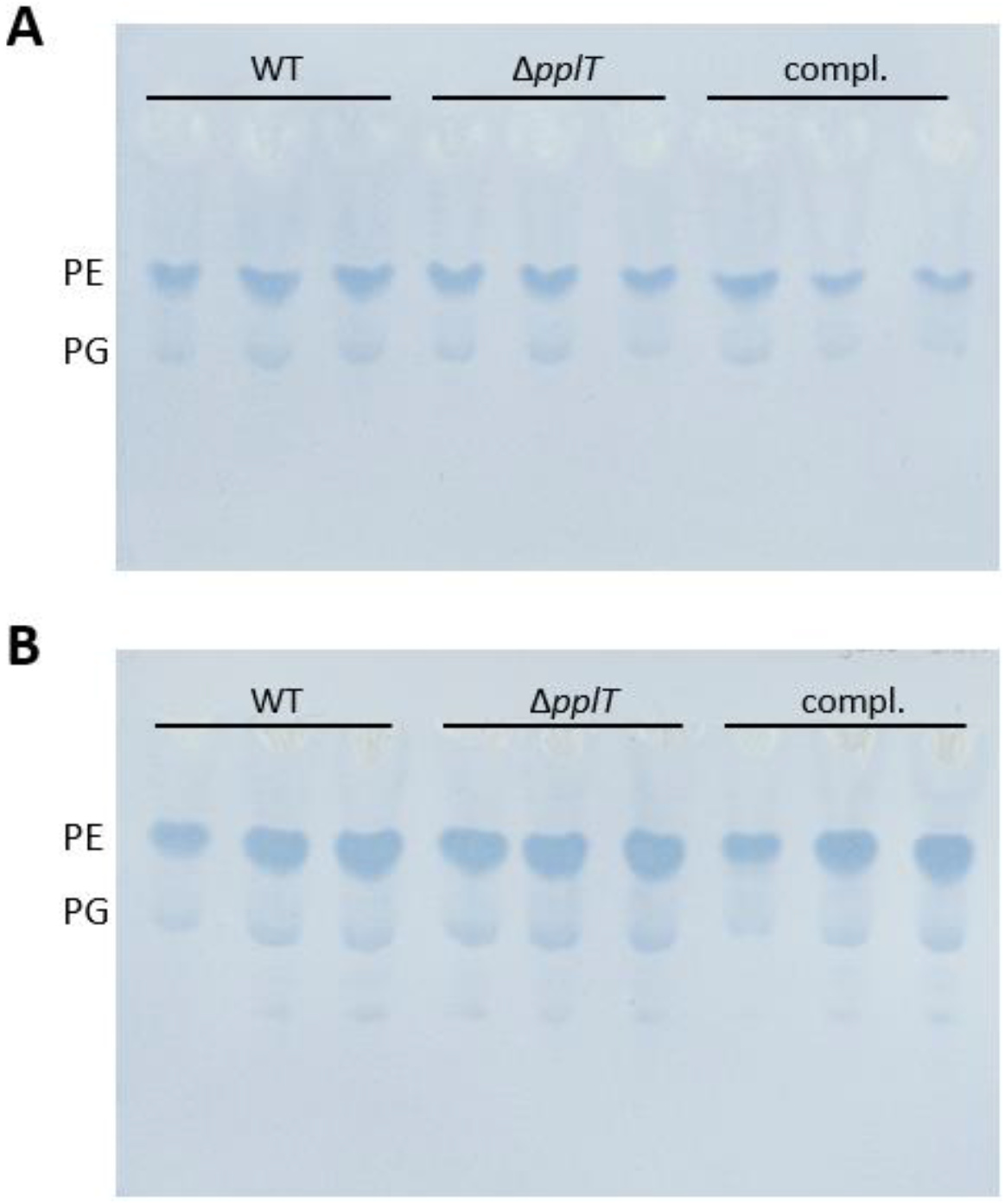
Impact of PplT on cell membrane lipid pattern. Detection of phospholipids of *E. coli* CFT073 WT, *pplT* mutant and complemented mutant at exponential (A) or stationary (B) growth phase. Total lipids were isolated, separated via thin-layer chromatography and stained using molybdenum blue spray reagent. PE and PG spots are indicated, concentrations of the minor lipids PS and CL are too low to be detectable. Three biological replicates are represented.

**Figure 5:**
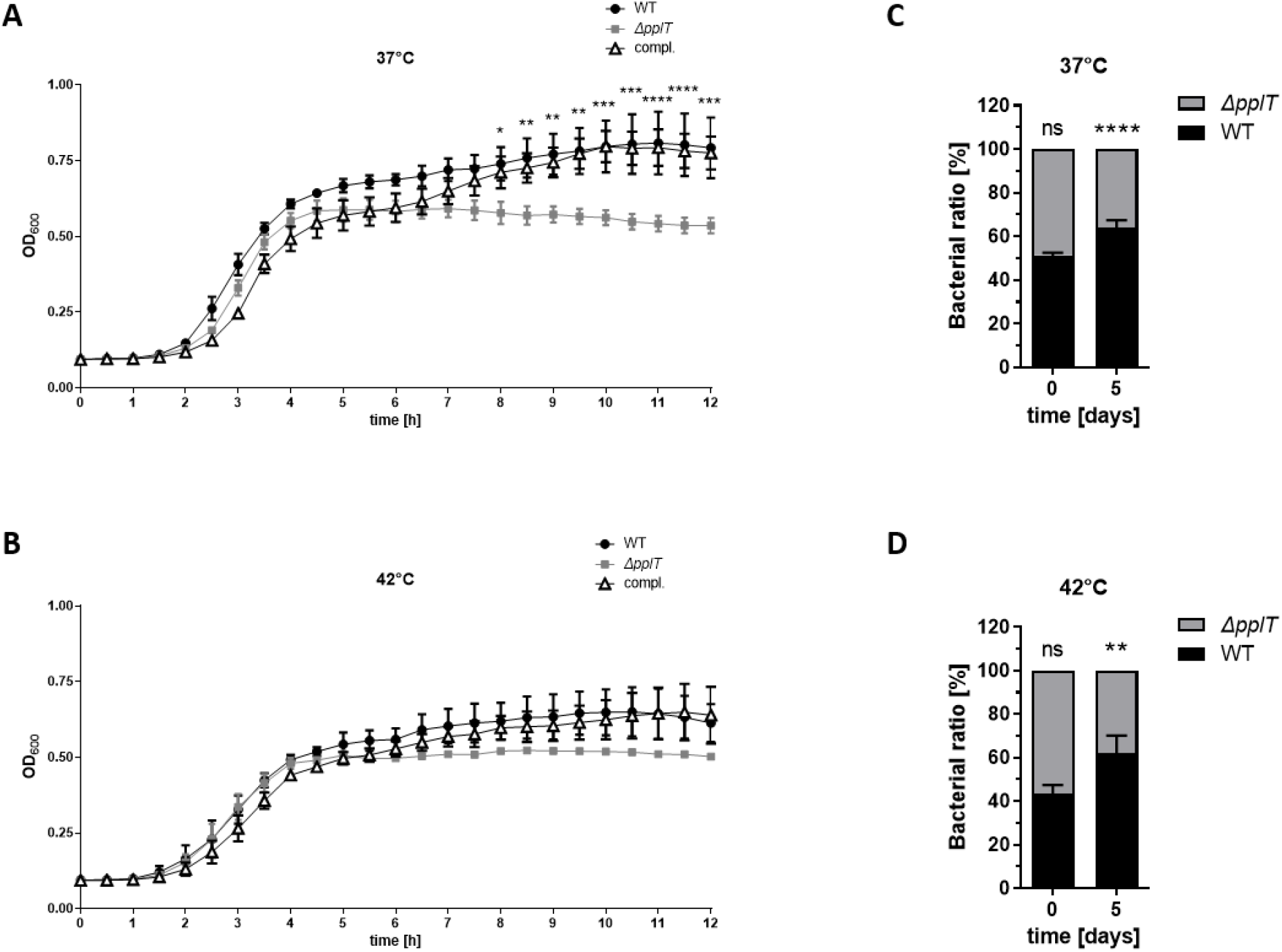
The *E. coli pplT* mutant exhibits a growth deficit compared to the WT. Growth curves of *E. coli* CFT073 WT, *pplT* mutant and complemented mutant at 37°C (A) or 42°C (B). Means of three biological replicates plus SEM are shown. The difference in growth between WT and *pplT* mutant was statistically analyzed by two-way ANOVA (*, P < 0.05; **, P < 0.01; ***, P < 0.0005; ****, P < 0.0001). Competition assay of WT and the *pplT* mutant over five days at 37°C (C) or 42°C (D). Means of three biological replicates with SD are shown. Data were statistically analyzed by two-way ANOVA and compared to the WT at the appropriate timepoint (ns, not significant; **, P < 0.01; ****, P < 0.0001).

## Discussion

Phospholipids are synthesized at the inner leaflet of the cytoplasmic membrane but they need to be translocated to the outer leaflet to form a stable bilayer [1]. Phospholipids can flip spontaneously but this process is rather inefficient [21]. Appropriate membrane homeostasis therefore depends on dedicated flippases that translocate lipids. While such transporters are known for eukaryotic cells, their presence and nature in prokaryotes has remained largely unclear. ABC transporters translocating the lipid-bound precursors of peptidoglycan or teichoic acids have been identified [22, 23]. However, these molecules are usually highly specific and are unlikely to have a major impact on phospholipid translocation. Nevertheless, the lipid-A translocating ABC transporter MsbA of *E. coli* has also a minor affinity for phospholipids [23]. The only dedicated bacterial phospholipid flippase described to date is the integral-membrane domain of MprF proteins, which connects the synthesis of aminoacyl phospholipids with their translocation [4, 5]. We show here that the flippase domain of MprF, named the PplT domain, can be found as a separate protein or combined in modular proteins with other domains of diverse function, which supports a general role of PplT in bacterial and archaeal cells.

The PplT protein modulates the fitness of *E. coli*. The observed effects were moderate, which may be based on the fact that the lipid A flippase MsbA has also some basic phospholipid translocation capacity [23] and may compensate for the lack of PplT to some degree. *msbA* is essential though [16, 24], which precludes its simultaneous inactivation with *pplT*. The residual capacity of MsbA cell wall precursor flippases to translocate phospholipids may also explain why several bacterial species do not seem to encode obvious PplT orthologs. Membrane function is strongly affected by temperature, which may explain the temperature-dependent impact of PplT in *E. coli*. It should be noted though that PplT domain proteins are also found in thermophilic microorganisms such as Thermotogae and archaeal extremophiles, suggesting that the translocation of certain lipids may require dedicated flippases even at high temperatures. The frequent combination of PplT with other enzymatic protein domains also suggests that certain modified lipids, potentially those with bulkier head groups, may be particularly dependent on cognate translocation machineries. Lipid translocation is a very dynamic process, which is difficult to monitor directly. Therefore, evidence for the flippase function of PplT proteins remains in part indirect. However, the fact that PplT interacts directly with most of the phospholipid-biosynthetic proteins of *E. coli* underscores its central role in membrane homeostasis. Likewise, MprF proteins have been found to interact with larger phospholipid-biosynthetic protein complexes in *S. aureus* [17, 18].

It remains unclear how wide or narrow the substrate specificities of PplT domain proteins may be. Since *E. coli* does not produce aminoacyl phospholipids, PplT may translocate some or all of the *E. coli* phospholipids PG, CL, PS, and PE. These molecules differ in the net charge of their head groups, which has been found to be a major determinant for substrate recognition by the PplT domain of the *R. tropici* MprF [7]. Although the PplT domain of the *S. aureus* MprF is linked to synthesis of the cationic phospholipid Lys-PG, MprF has been found to also translocate the zwitterionic phospholipid Ala-PG [25], indicating that PplT proteins may have broader substrate specificities. It should be noted that most prokaryotic genomes with a PplT homolog contain only one PplT protein and only a minority encodes two (19.89%) or more (1.60%) of them, suggesting that one flippase may usually be sufficient for most of the membrane-forming phospholipids. Archaea have different phospholipids compared to bacterial ones, with monolayer-membrane forming isoprenoid lipids [27]. The broad presence of PplT proteins in Archaea suggest that these proteins may also be able to flip such lipids, which are much bulkier than bacterial phospholipids and contain two head groups, one at each end. It remains unclear if the PplT domain may only facilitate the exchange of phospholipids between the inner and outer leaflet of cytoplasmic membranes or if it can translocate phospholipids in an energy-dependent fashion to generate asymmetric lipid patterns. Some of the transmembrane domains of MprF seem to be related to the proton-motif force-dependent major facilitator protein superfamily [28] and some studies found Lys-PG to be unevenly distributed between the two membrane leaflets [4], suggesting that PplT may use energy to translocate its substrate lipids.

Membrane homeostasis is an important fitness factor for all kinds of organisms. PplT proteins may therefore become attractive targets for future anti-fitness drugs to combat major human pathogens. In support of such a strategy, monoclonal antibodies directed against extracellular portions of the PplT domain of the *S. aureus* MprF have recently been shown by inhibiting the Lys-PG exposure at the outer leaflet of the membrane and therefore sensitize *S. aureus* to cationic antibiotics [9].

## Materials and Methods

### Occurrence and architecture of PpIT in bacterial and archaeal taxa

In order to determine the distribution of PpIT across the taxonomy a blastp search within NCBI’s non-redundant protein database [29] was performed using the PplT/YbhN protein sequence of *E. coli* (old: NP_752801.1; new: WP_000045478.1) as a query and an E-value cutoff of 0.05. Blast hits originating from the superkingdoms Bacteria and Archaea were investigated separately. For each, the subtree of the NCBI taxonomy rooted at the respective superkingdom was thinned out for those taxa that possess a PpIT homolog. The BLAST search and subsequent taxonomic visualization was performed with BLASTphylo [https://github.com/Integrative-Transcriptomics/BLASTphylo]. We used NCBI’s Conserved Domain Architecture Retrieval Tool (CDART) [15] to identify different domain architectures that include a PpIT domain. We used MicrobesOnline[30] to identify *pplT*’s operon structure in other Enterobacteriaceae.

### Bacterial strains, mutagenesis, and maintenance

*E. coli* UPEC strain CFT073 (DSM103538) (Tab. S1) was used for *pplT* mutagenesis and analysis [31]. *pplT* was deleted in the chromosome of CFT073 as described previously [32]. Briefly, *E. coli* CFT073 was transformed with helper plasmid pKD46 encoding the Lambda-Red recombinase (Tab. S2) and incubated at 30°C on LB agar supplemented with ampicillin. The kanamycin resistance cassette with chromosomal DNA regions flanking *pplT* was amplified by PCR with primers pplT_KO_5 and pplt_KO_6 (Tab. S3) using pKD13 as template (Tab. S2). After arabinose-induced Lambda Red expression in *E. coli* CFT073 containing the helper plasmid, the PCR fragment was transferred by electroporation and cells were plated on kanamycin-containing LB agar for primary selection. After verification of the chromosomal *pplT* deletion by PCR (Tab. S3) and sequencing, mutants were streaked to single colonies on LB agar without antibiotics at 37°C twice for curing of the temperature-sensitive helper plasmid. The loss of the helper plasmid pKD46 was confirmed by PCR (Tab. S3). The *pplT* deletion mutant was complemented by a *pplT* gene amplified from *E. coli* K12 and cloned in the *E. coli*/*S. aureus* shuttle vector pRB474 [33] (Tab. S2). All bacterial strains were grown in LB medium (Carl Roth) with appropriate antibiotics unless otherwise noted (Tab. S1).

### Bacterial two-hybrid assay

To identify potential interaction partners of PplT, the commercially available bacterial two-hybrid kit (BATCH kit, Euromedex) was used [8, 19, 20] and the proteins to be studied were cloned as described previously [8]. The *pplT* gene was cloned in the high-copy number plasmid pUT18C, the potential interaction partners in the low copy-number plasmid pKT25 leading to C-terminal or N-terminal fusion with the adenylate cyclase fragments T25 or T18 of *Bordetella pertussis,* respectively (Tab. S2). The constructs were used to co-transform chemically competent *E. coli* BTH101. The resulting transformants were tested in a 96-well format for β-galactosidase activity, to quantify the protein-protein interaction as described previously [8] with slight modifications. Briefly, bacterial cells were grown at 30°C overnight in LB, supplemented with 0.5 mM isopropyl-β-D-thiogalactopyranoside (IPTG), 25 μg/ml kanamycin and 100 μg/ml ampicillin. 100 μl of the overnight cultures were used to measure the optical density at 600 nm (OD_600_) in a 96-well microtiter plate (F-bottom, Falcon), another 100 μl were transferred into a deep 96-well plate (U-bottom, Sarstedt) and mixed with 1 ml buffer Z (60 mM Na_2_HPO_4_, 40 mM NaH_2_PO_4_, 10 mM KCl, 1 mM MgSO_4_, 50 mM β-mercaptoethanol). To lyse the cells, 40 μl sodium dodecyl sulfate (0.1%) and 80 μl chloroform were added and mixed vigorously. After incubation at room temperature to allow phase separation, 100 μl of the aqueous phase were transferred into a 96-well microtiter plate (F-bottom, Falcon) and mixed with 20 μl o-nitrophenyl-β-D-galactopyranoside (4 mg/ml) to start the enzymatic reaction. The reaction was measured continuously for 1 h in a CLARIOStar plate reader (BMG Labtech) at OD_420_ and OD_550_. The β-galactosidase activity in Miller Units was calculated by the formula: 1000*((OD_420_-(1.75*OD_550_))/(t*v*OD_600_)), where t is reaction time in minutes and v the reaction volume in milliliter.

### Isolation of polar lipids

Polar lipids were extracted from bacteria grown to exponential or stationary phase using the Bligh-Dyer procedure [34]. Briefly, phospholipids were extracted by a mixture of sodium acetate buffer (20mM, pH 4.8), chloroform, and methanol (1:1:1 [by volume]), vacuum dried, and dissolved in chloroform-methanol (2:1 [by volume]). For detection of phospholipid patterns, appropriate amounts of polar lipid extracts were spotted onto silica gel high-performance thin-layer chromatography (HPTLC) plates (silica gel 60 F_254_, Merck) using a Linomat 5 sample application unit (CAMAG) and developed with chloroform:methanol:water (65:25:4 [by volume]) in an automatic developing chamber ADC 2 (CAMAG). Phospholipids were selectively stained with molybdenum blue spray reagent (1.3% in 4.2 M sulfuric acid, Sigma-Aldrich).

### Bacterial growth assay

Temperature-dependent growth differences were analyzed in 96-well microtiter plate format. To this end, overnight cultures grown in LB with appropriate antibiotics were adjusted to OD_600_ 0.05 in fresh LB and 100 μl of the cell suspensions were transferred to a 96-well microtiter plate (F-bottom, Falcon) and growth was measured at OD_600_ in 30 min intervals at either 37°C or 42°C in an Epoch2 plate reader (BioTek).

To investigate growth differences of WT and *pplT* mutant in cocultivation, overnight cultures were adjusted to the same colony forming unit in fresh LB and used to inoculate a coculture, which was incubated in a shaking incubator at either 37°C or 42°C. Cells were passaged every 24 h in fresh LB for five days. To measure the numbers of live cells of the two competing strains, appropriate dilutions of the suspensions were spotted on LB agar plates with or without kanamycin. After incubation of the agar plates at appropriate temperatures, colonies were counted and ratios were calculated.

### Statistics

Statistical analyses were performed with the Prism 8.4.2 package (GraphPad Software) and Group differences were analyzed for significance with one-way or two-way ANOVA. A *P* value of ≤0.05 was considered statistically significant.

## Acknowledgments

We thank Anne Berscheid for providing plasmids pKD13 and pKD46, Cordula Gekeler and Ulrike Redel for technical support and Jennifer Müller for her contributions to the BLASTphylo project. This work was financed by Grants from Deutsche Forschungsgemeinschaft (DFG) SFB766 (to A.P.) and TRR261, project ID 398967434 (to K.N. and A.P), the German Center of Infection Research (DZIF) (to A.P. and C.S.). C.S. was also supported by the intramural Experimental Medicine program of the Medical Faculty at the University of Tübingen. The authors acknowledge project support (C.S.) and infrastructural support (C.S., K.N., A.P.) by the DFG Cluster of Excellence EXC2124 CMFI, project ID 390838134.

## Supporting Information

**Table S1:** Bacterial strains.

| Strain | Characteristics |
| --- | --- |
| *E. coli* CFT073 | Uropathogenic wild type strain [31] |
| *E. coli* CFT073 $\Delta ppIT$ | *ppIT* chromosomal deletion mutant, *ppIT* gene was replaced by kanamycin resistance cassette, Kan<sup>R</sup>. This study. |
| *E. coli* CFT073 $\Delta ppIT$ pRB*ppIT* | *ppIT* deletion mutant complemented by plasmid-encoded *ppIT* version. This study. |

**Table S2:** Plasmids.

| Plasmid | Characteristics |
| --- | --- |
| pRB474*ppIT* | *ppIT* gene cloned in *E. coli*/*S. aureus* shuttle vector pRB474 [33], Amp<sup>R</sup>. |
| pKD13 | Template for Kanamycin resistance cassette. Amp<sup>R</sup>, Kan<sup>R</sup> [32]. |
| pKD46 | Lambda Red helper plasmid, Lambda Red expression inducible by L-arabinose. Amp<sup>R</sup> [32]. |
| pKT25-*clsA* | *clsA* gene encoded on low copy number plasmid pKT25 (Euromedex), fused to the C-terminal end of the adenylate cyclase fragment T25. This study. |
| pKT25-*clsB* | *clsB* gene encoded on low copy number plasmid pKT25 (Euromedex), fused to the C-terminal end of the adenylate cyclase fragment T25. This study. |
| pKT25-*clsC* | *clsC* gene encoded on low copy number plasmid pKT25 (Euromedex), fused to the C-terminal end of the adenylate cyclase fragment T25. This study. |
| pKT25-*ybhP* | *ybhP* gene encoded on low copy number plasmid pKT25 (Euromedex), fused to the C-terminal end of the adenylate cyclase fragment T25. This study. |
| pKT25-*pssA* | *pssA* gene encoded on low copy number plasmid pKT25 (Euromedex), fused to the C-terminal end of the adenylate cyclase fragment T25. This study. |
| pKT25-*psd* | *psd* gene encoded on low copy number plasmid pKT25 (Euromedex), fused to the C-terminal end of the adenylate cyclase fragment T25. This study. |
| pKT25-*pgsA* | *pgsA* gene encoded on low copy number plasmid pKT25 (Euromedex), fused to the C-terminal end of the adenylate cyclase fragment T25. This study. |
| pKT25-*pgpA* | *pgpA* gene encoded on low copy number plasmid pKT25 (Euromedex), fused to the C-terminal end of the adenylate cyclase fragment T25. This study. |
| pKT25-*pgpB* | *pgpB* gene encoded on low copy number plasmid pKT25 (Euromedex), fused to the C-terminal end of the adenylate cyclase fragment T25. This study. |
| pKT25-*pgpC* | *pgpC* gene encoded on low copy number plasmid pKT25 (Euromedex), fused to the C-terminal end of the adenylate cyclase fragment T25. This study. |
| pKT25-*zip* | Leucine zipper of GCN4 fused to the C-terminal end of the adenylate cyclase fragment T25. BATCH system kit, Euromedex. |
| pUT18C-*ppIT* | *ppIT* gene encoded on high copy number plasmid pUT18C (Euromedex), fused to the C-terminal end of the adenylate cyclase fragment T18. This study. |
| pUT18C-*zip* | Leucine zipper of GCN4 fused to the C-terminal end of the adenylate cyclase fragment T18. BATCH system kit, Euromedex. |

**Table S3:**
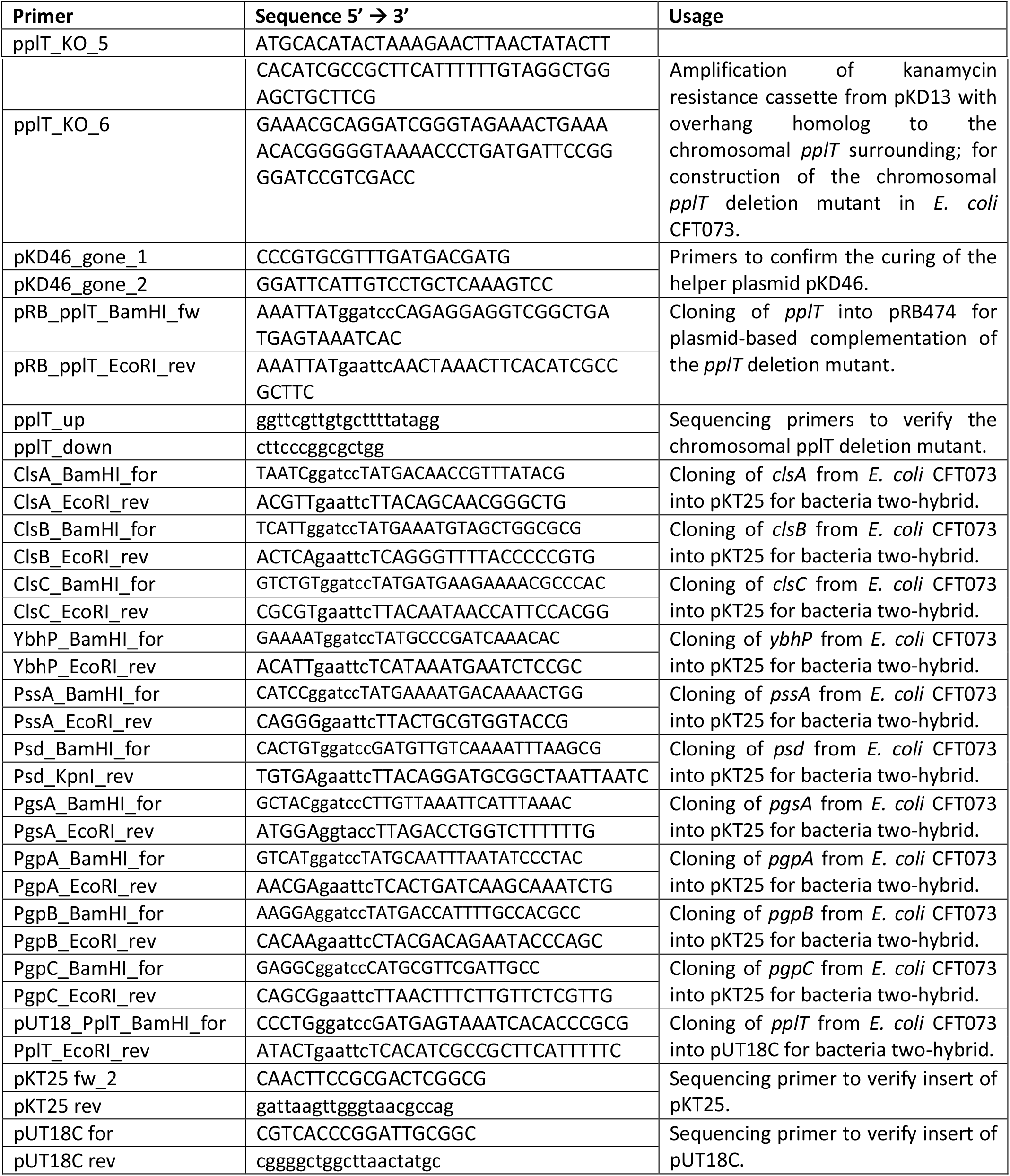
Primers.

**Figure S1:**
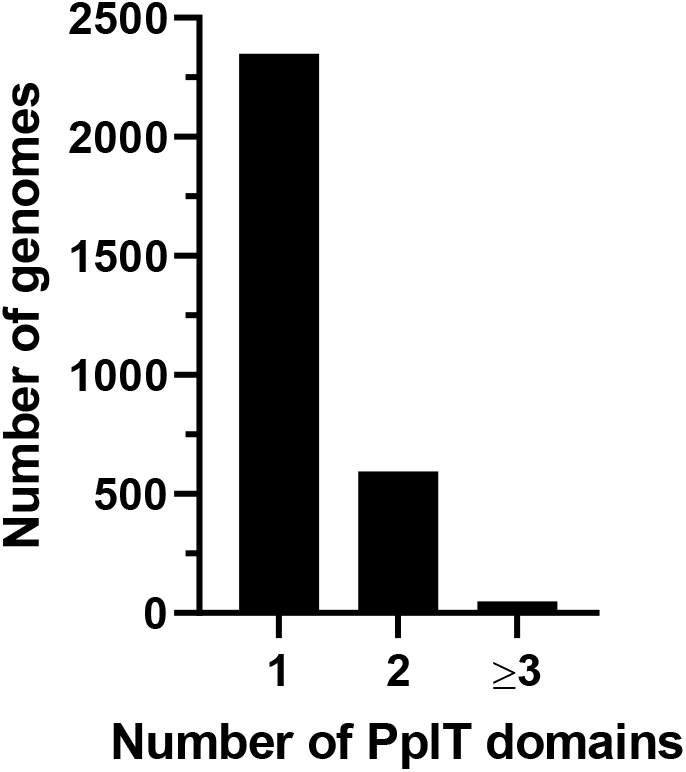
Most PplT containing genomes of prokaryotes harbor only one homolog. Number of genomes with one, two or more genes encoding PplT domains are shown.

